# Biofilm dispersion in *Enterococcus faecalis* is mediated by nutrient step-change and intra-species signaling

**DOI:** 10.64898/2026.05.20.724677

**Authors:** Nermin Mohamed, David M. Lam, Marian Abdikarin, Raheema Mohammed-Abraham, David G Davies, Laura C Cook, Peter T. McKenney

**Affiliations:** Binghamton Biofilm Research Center, Department of Biological Sciences, Binghamton University, SUNY, Binghamton NY 13902 USA; Department of Immunology and Microbial Disease, Albany Medical College, Albany NY 12208 USA

## Abstract

*Enterococcus faecalis* is a Gram-positive intestinal commensal and opportunistic pathogen capable of causing serious infections, including urinary tract infections, endocarditis, and wound infections. A major contributor to its persistence during infection is the ability to form biofilms on host tissues and medical devices. Biofilm cells have higher phenotypic tolerance to antimicrobial treatment than planktonic bacteria. While mechanisms governing biofilm assembly in *E. faecalis* have been widely studied, the processes that regulate biofilm dispersion, the final stage of the biofilm life cycle, remain poorly understood. In this study, we found that dispersion is triggered by a tenfold step-change increase in nutrient availability and by cell free supernatant (CFS) of *E. faecalis* OG1RF cultures. Cells released from biofilms regain sensitivity to antibiotics similar to planktonic cells but maintain a high potential for adherence. We characterized the glycosyltransferase *epaOX*, which contributes to the structure of the enterococcal polysaccharide antigen as necessary for nutrient step-change induced dispersion, CFS induced dispersion, and adhesion of dispersed cells. Supplementation of *epaOX* mutant CFS with galactose and N-acetylgalactosamine was sufficient to restore CFS induced dispersion. Together these data suggest that dispersion in OG1RF occurs with fast kinetics, affects antibiotic sensitivity and is regulated in part by known virulence factors.

**Importance:** *E. faecalis* causes difficult to treat infections at numerous body sites in human patients. *E. faecalis* biofilms are adherent populations that require high levels of antibiotics for treatment. Biofilms undergo a disassembly process named dispersion that allows individual cells to leave the biofilm and colonize new locations. Dispersed cells in other species are killed by lower amounts of antibiotics than biofilm cells. Here we showed that dispersion occurs in *E. faecalis* and lowers the level of antibiotics needed to kill dispersed cells. Dispersion triggers could be used in the future to design treatments that increase the effectiveness of antibiotics.

## Introduction

One of the most significant biofilm-forming organisms that cause nosocomial infections is *Enterococcus faecalis*. Biofilm formation in *E. faecalis* plays a critical role in the pathogenesis and persistence of many challenging-to-treat infections, such as periodontitis, endocarditis and urinary tract infections (Arias and Murray 2012). As in other species of bacteria, biofilms of *E. faecalis* have increased phenotypic tolerance to antibiotics used in treatment (Frank *et al*. 2015). Combined with increasing genetic resistance to antibiotics, high levels of biofilm antibiotic tolerance may lead to treatment failures in *E. faecalis* infections. Biofilm development in *E. faecalis* is a multi-step process of attachment, microcolony formation, and maturation (Willett and Dunny 2024; Ch’ng et al. 2019). In other systems, biofilm dispersion or disassembly has been characterized (Rumbaugh and Sauer 2020). Several dispersion cues have been shown to trigger the removal of up to 80% of biofilm biomass in as little as 1 hour in *Pseudomonas aeruginosa* and other species (Barraud et al. 2006; Davies and Marques 2009; K. Sauer et al. 2004). Dispersion is important to understand mechanistically because cells released from the biofilm quickly lose their high levels of phenotypic antibiotic tolerance (Chambers, Cherny, and Sauer 2017; Dingemans et al. 2018; Goodwine et al. 2019). This suggests that dispersion, if it could be triggered without stimulating systemic infection, could provide a promising strategy for increasing the efficacy of antibiotics in treating biofilm infections.

Biofilm dispersion in *E. faecalis* largely has not been characterized mechanistically. Some reports have characterized the effects of various treatments, such as berberine, D-amino acids, and glycoside hydrolase enzymes, on mature *E. faecalis* biofilms and reported effects on disassembly or dispersion (Chen et al. 2016; Fleming et al. 2020; Rosen et al. 2016). These assays generally were designed to test potential inhibitory agents on biofilms and often used changes in biomass as a readout after overnight exposure. In other systems the kinetics of dispersion is fast and can be measured in minutes using continuous flow (K. Sauer et al. 2004) or batch culture systems (Davies and Marques 2009), or time-lapse imaging (Vidakovic et al. 2023). Here we sought to develop an assay that could be used to describe dispersion in *E. faecalis* on a similar time scale and interrogate its genetic regulation.

In this study, we characterize biofilm dispersion in the model organism *E. faecalis* OG1RF. We present evidence that biofilm dispersion can be induced by a sudden step-change in nutrient availability. We also show that a step-change in nutrients results in the loss of antibiotic tolerance of the released cells to multiple clinically relevant antibiotics, regardless of the mechanism of action. We show that dispersion can be induced by cell free supernatant and that the gene *epaOX* is necessary to induce dispersion of wild type biofilms. Taken together these data suggest that *E. faecalis* integrates multiple environmental cues into decisions to remain in a biofilm.

## Results

Dispersion is classified as ‘native dispersion’, if it occurs as a phenomenon triggered by cues from within the community (Davies and Marques 2009). To determine if OG1RF undergoes native dispersion, we grew biofilms in multiwall plates for 8 days while replacing the spent medium with fresh medium every 24 hours. If native dispersion occurred, we would expect to see a peak in recovered CFUs in the bulk liquid after overnight growth. However, OG1RF formed stable biofilms from day 2 to day 8 under these conditions (Fig S1) and we did not detect obvious native dispersion in these batch culture conditions. We observed a continuous increase in biofilm biomass by crystal violet staining (Fig S1C), but no change in total biofilm CFUs over time from day 2 to day 8 (Fig S1D-E). We observed a not significant trend towards decreased biofilm biomass and thickness between days 4 and 8 using confocal laser scanning microscopy (Figure S2). Based on these data, we chose day 4 as the experimental time point for induced biofilm dispersion assays.

### Nutrient Step-Change Induces Biofilm Dispersion in *E. faecalis* OG1RF

In *Pseudomonas aeruginosa*, a sudden increase in carbon availability has been shown to trigger dispersion, resulting in an 80% reduction in biofilm biomass (K. Sauer et al. 2004). We investigated whether a similar nutrient-responsive dispersal mechanism exists in OG1RF. OG1RF biofilms were cultivated under nutrient-limited conditions (10% TSB) for 4 days in multi-well plates with daily medium change. On day 4, the overnight bulk liquid was removed, and biofilms were gently rinsed and incubated for 1 h in fresh 10% TSB to standardize baseline nutrient exposure. To induce dispersion, a step-change in nutrients was then introduced by replacing the 10% TSB with 100% TSB for 1 h. Following treatment, cells released from the biofilm into the bulk liquid and remaining attached biofilms were collected separately and enumerated as colony-forming units (CFUs). We observed a strong trend towards an increase in CFUs present in the liquid (p = 0.0510) and no change in CFUs of the attached biofilm (Fig 1A). When we plot these data as percentage of released cells, there was a significant change of approximately 6-fold (Fig 1B). We confirmed that growth and division of OG1RF does not account for this change by comparison with the growth of mid-log OG1RF under these conditions (Fig S3).

**Figure 1.**
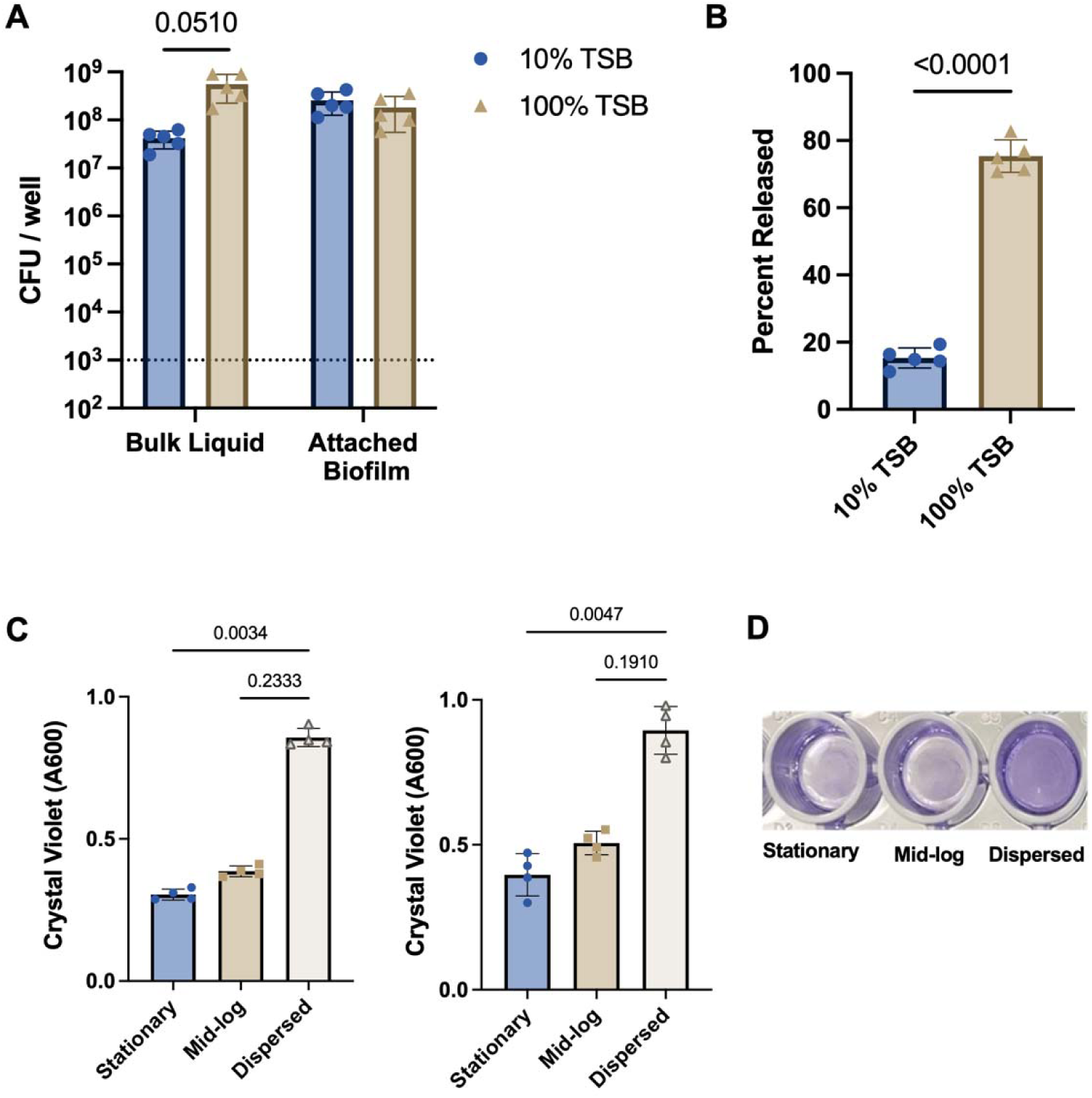
Nutrient step-change induces biofilm dispersion in *E. faecalis* OG1RF. (A-B) Nutrient-induced dispersion of 4-day mature OG1RF biofilms. Biofilms grown under nutrient-limited conditions (10% TSB) were exposed to a 10-fold nutrient upshift (100% TSB) for 1 hour followed by selective plating of the bulk liquid and the remaining attached biofilm, n = 5 biological replicates. A) CFU counts, unpaired t-test with Welch’s correction and Holm-Sidak multiple comparisons test. B) Data from A represented as percentage of cells into the bulk liquid, unpaired t-test with Welch’s correction. **C)** Attachment capacity of dispersed cells compared with planktonic and stationary phase populations. Dispersed cells were added to a 96-well plate an allowed to attach for 2 hours (left) or 24 hours (right) before crystal violet staining. n = 4 biological replicates, Kruskal-Wallace one-way ANOVA with Dunn’s test. D) Representative image of crystal violet staining from 24-hour attachment assay.

To determine if dispersed cells retain the high adherence of biofilm cells, we measured cell adherence relative to mid-log and stationary phase OG1RF. Using a 96-well plate–based attachment assay, we observed that dispersed cells from OG1RF biofilms had significantly higher attachment compared with both exponential phase and stationary phase planktonic cells at 2 and 24 hours post inoculation (Fig 1C-D). To determine whether nutrient step-change was accompanied by structural remodeling of the biofilm, we performed confocal laser scanning microscopy. Biofilms exposed to 100% TSB exhibited significantly reduced biofilm biomass and a trend toward reduced surface area when compared with 10% TSB control biofilms indicating disassembly had occurred in these conditions (Fig 2). Taken together, these data suggest that cells are released from OG1RF biofilms following a nutrient step change. Dispersed cells maintain a high capacity for adhesion compared to mid-log planktonic cells which may facilitate rapid recolonization of new surfaces.

**Figure 2.**
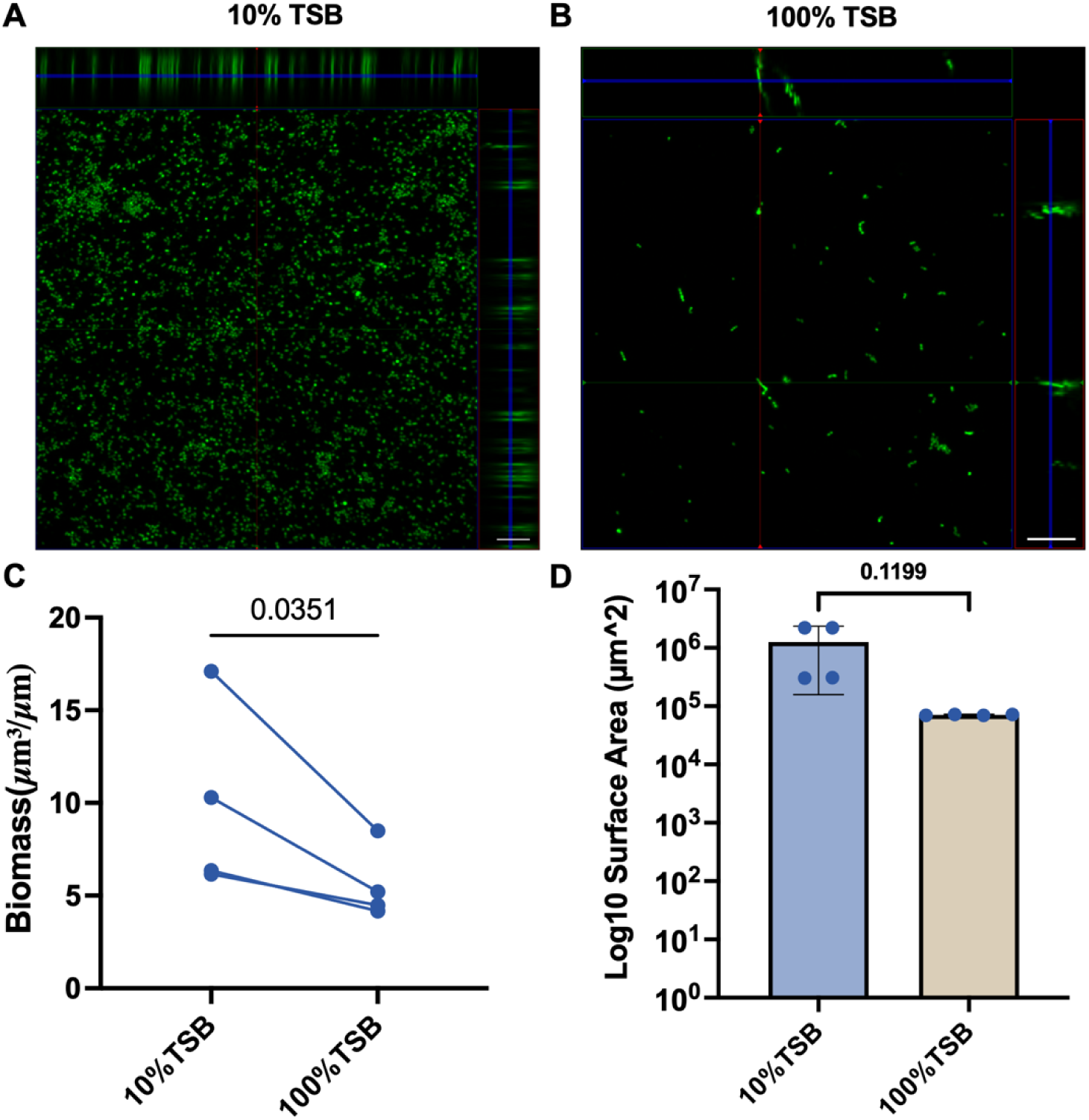
Confocal laser scanning microscopy (CLSM) analysis of biofilm architecture following nutrient step-change. Representative images of day 4 biofilms grown in 24-well plates under A) uninduced (10% TSB for 1 hour) or B) dispersion-induced (100% TSB for 1 hour, right) conditions. Scale bar = 100 µm. C) Quantitative analysis of biomass, D) surface area using COMSTAT2. Each data point represents the mean of 3–4 technical replicate wells, (n=4 biological replicates), paired two-tailed t-test.

### Dispersed cells lose phenotypic tolerance to antibiotics

In *P. aeruginosa*, dispersed cells are not identical to planktonic cells grown in liquid culture and possess a distinct phenotype (Sauer et al. 2002). Free-living mid-log planktonic cells are highly susceptible to antimicrobial agents compared to bacteria living in a biofilm. This phenotypic antibiotic tolerance raises the concentration of antibiotics required to kill cells 10 to 1,000-fold (Rumbaugh and Sauer 2020; Frank et al. 2015). We tested the antibiotic susceptibility of wild-type biofilm cells dispersed by nutrient step-change to a panel of clinically relevant antibiotics: Ampicillin, Vancomycin, Daptomycin, and Linezolid. Antibiotic concentrations were evaluated with planktonic mid-log cells to determine the lowest concentration that causes at least a 1-log decrease in CFUs following 1 hour of exposure (Fig S4). Across all antibiotics tested, cells in the bulk liquid following nutrient step-change (100% TSB) exhibited no difference in susceptibility compared to mid-log planktonic cells. For example, Vancomycin treatment resulted in approximately a 1.5-log reduction in both mid-log planktonic cells and released biofilm cells in the bulk liquid, whereas intact attached biofilm cells showed minimal killing (∼0.2-log reduction) (Fig 3A). Similar trends were observed with Linezolid, Ampicillin, and Daptomycin, where released cells in the bulk liquid showed levels of susceptibility similar to mid-log planktonic cells (Fig 3B-D). Together, these findings demonstrate that nutrient-induced dispersion results in a physiological shift associated with loss of biofilm-associated antibiotic tolerance, independent of antibiotic class or mechanism of action.

**Figure 3.**
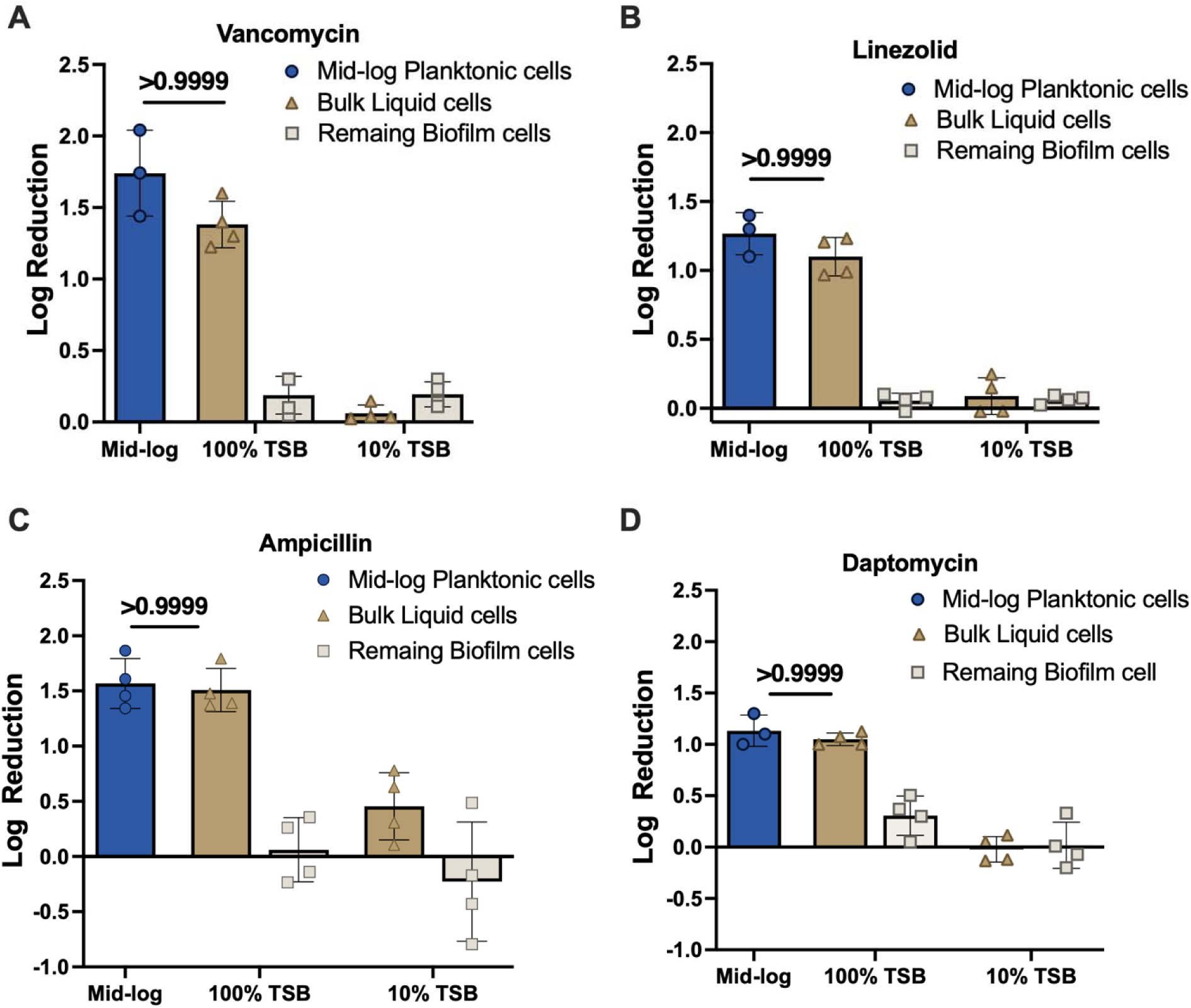
Nutrient step-change eliminates phenotypic antibiotics tolerance in released cells independent of mechanism of action. Four-day mature OG1RF biofilms were subjected to a nutrient step-change or control for 1 hour. Released cells in the bulk liquid and attached biofilms were collected and exposed to the indicated antibiotic. Mid-log planktonic OG1RF in grown in 10% TSB were exposed to the indicated antibiotic as a control. Data are plotted as log reduction relative to untreated cells of the same fraction. Antibiotic concentrations were chosen based on titration (Figure S2) A) Vancomycin 50 μg/mL, B) Linezolid 200 μg/mL, C) Ampicillin 50 μg/mL, D) Daptomycin 50 μg/mL + 75 μg/mL CaCl_2_. Each data point represents the mean of 3–4 technical replicate wells from 4 independent experiments (n=3-4 biological replicates). Statistics: Kruskal-Wallace one-way ANOVA with Dunn’s correction versus Mid-log.

### Dispersion is induced by OG1RF cell-free supernatant

To determine if OG1RF itself produces extracellular factors capable of promoting biofilm dispersal, we examined the effect of cell-free supernatants (CFS) for their capacity to induce dispersion. CFS obtained from 24-hour, and 72-hour cultures grown in 10% TSB resulted in a significant increase in the percentage of cells released from the biofilm (Fig 4A). To investigate the nature of the dispersal-inducing factor(s), the CFS was subjected to treatments that target protein stability. Heat treatment of the CFS at 95 °C for 30 min led to a significant decrease in the percentage of released cells (Fig 4B), indicating that the dispersal activity is partially heat-labile. Similarly, treatment of the CFS with Proteinase K resulted in a trend towards reduction in dispersal activity (p = 0.064) (Fig 4C). Together, these findings suggest that proteinaceous components present in the extracellular milieu may contribute to the observed biofilm dispersal activity, although additional non-protein factors may also be involved.

**Figure 4.**
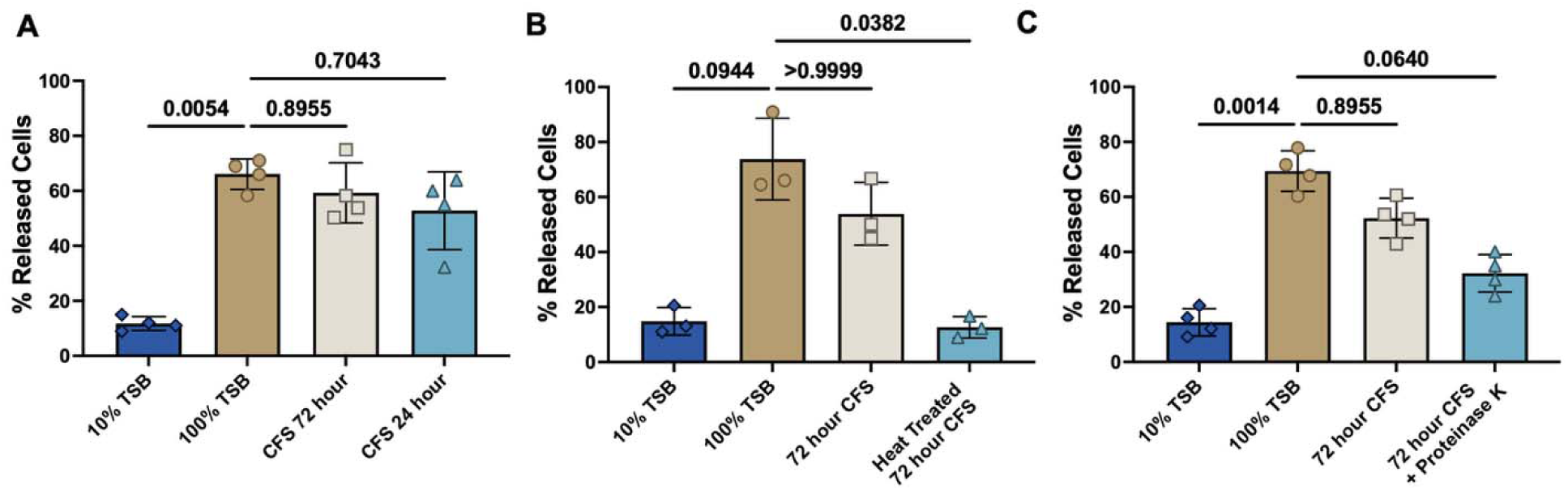
Dispersion is induced by cell free supernatant. 4-day biofilms were exposed to the indicated treatment. Bulk liquid and attached biofilms were collected and CFUs were quantified. Data are presented as percentage of released cells. Data represent 3-4 independent experiments. Statistics: Kruskal-Wallace one-way ANOVA followed by Dunn’s multiple comparisons test. **(A)** CFS = Cell free supernatant from a wild type OG1RF culture in 10% TSB taken 72 or 24 hours post inoculation. **(B)** CFS subjected to heat treatment (95°C for 30 min). **(C)** CFS subjected to Proteinase K treatment for 1 hour followed by heat inactivation.

### EpaOX is necessary for CFS-induced dispersion

One gene that has the potential to affect dispersion and antibiotic tolerance in biofilms is the glycosyl transferase *epaOX* (Dale et al. 2015). Mutants in *epaOX* had decreased biofilm growth in the presence of antibiotics (Dale et al. 2015) and increased detachment of biofilms following treatment with daptomycin (Dale et al. 2017). We tested a transposon insertion in *epaOX* (Fig S5) for effects on induction of biofilm dispersion. Biofilms of *epaOX::Tn* released a similar percentage of cells in response to nutrient step change to those treated with 10% TSB, suggesting that *epaOX* is necessary for nutrient-induced dispersion within established biofilms (Fig 5A). We applied cell-free supernatant (CFS) derived from *epaOX::Tn* liquid culture applied to wild-type biofilms and observed a significant reduction in the percentage of released cells (Fig 5B). These results suggest that EpaOX may be necessary for nutrient step-change induced dispersion and for producing extracellular secreted factors that promote biofilm dispersion induced by CFS. We also found that *epaOX::Tn* dispersed cells did not have the high levels of attachment relative to planktonic cells observed for wild type OG1RF (Fig 5C). These data suggest that EpaOX and potentially the Epa polysaccharides mediate dispersion and re-attachment of biofilm cells.

**Figure 5.**
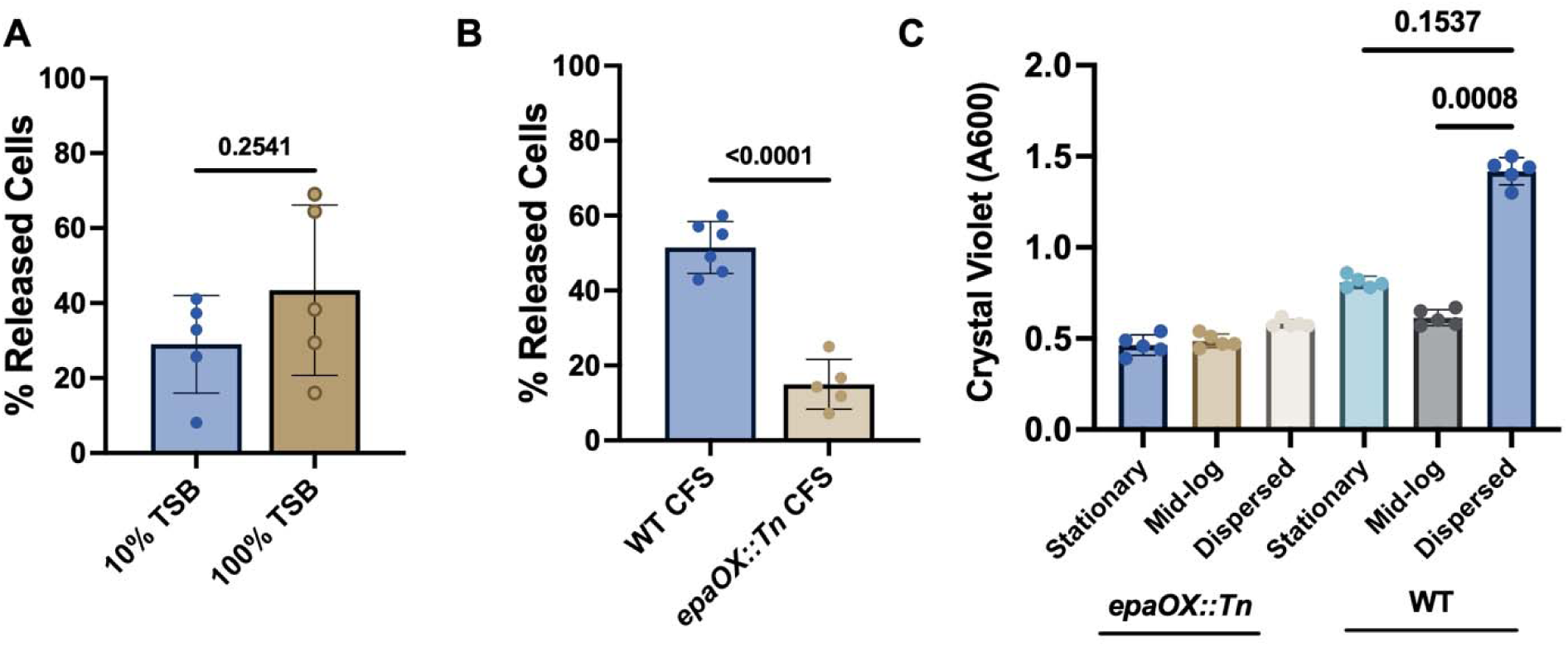
EpaOX is necessary for dispersion in response to cell free spent medium and reattachment of dispersed cells. **(A)** Four-day mature biofilms of the *epaOX::Tn* mutant were subjected to nutrient step-change (10% to 100% TSB) or control for 1 hour followed by harvest of the bulk liquid and attached biofilm and quantification of CFUs. Data are presented as percentage of released cells. Data represent 5 independent experiments. **(B)** Cell-free supernatant (CFS) collected from 72-hour cultures of *epaOX::Tn* or wild-type OG1RF was applied to 4-day wild-type biofilms for 1 hour followed by harvest of the bulk liquid and attached biofilm and quantification of CFUs. Data are presented as percentage of released cells. Data represent 5 independent experiments. Statistics: unpaired Welch’s t-test. **(C)** 2-hour attachment of dispersed cells versus stationary phase and mid-log cultures of the indicated genotype. Data represent 5 independent experiments. Statistics: Kruskal-Wallace One-way ANOVA with Dunn’s test versus dispersed.

The gene encoding EpaOX is located in the enterococcal polysaccharide antigen (*epa*) gene cluster (Dale et al. 2015). Polysaccharides extracted from *epaOX* mutant strains are altered compared to wild type (Dale et al. 2015; Korir, Dale, and Dunny 2019), with a reduction in rhamnopolymer bands on a non-denaturing polyacrylamide gel. We confirmed that polysaccharide production is disrupted in extracts from the *epaOX::Tn* strain (Figure S5D); however, polysaccharide extracts themselves did not induce biofilm dispersion (Figure S6). Because carbohydrate-enriched extracts did not induce biofilm dispersion, we next examined whether individual monosaccharides that constitute these polysaccharides could influence the dispersal response. The enterococcal polysaccharide antigen (Epa) is composed of repeating sugars including galactose and N-acetylgalactosamine (GalNAc) (Davis et al. 2025). The abundance of monosaccharides galactose and GalNAc were reduced in previous mass spectrometry analysis of *epaOX* mutants (Dale et al. 2017). We tested the effect of these sugars added to CFS on mature biofilms for the ability to induce dispersion. Addition of GalNAc or galactose alone, or in combination to wild type CFS did not significantly affect dispersion of wild-type biofilms (Fig 6). When GalNAc or galactose was added singly or in combination to *epaOX::Tn* CFS it significantly restored induction of dispersion. Taken together, these data suggest that EpaOX influences dispersion via the production of free GalNAc and galactose. Furthermore, GalNAc and galactose act in part as small molecule triggers of biofilm dispersion.

**Figure 6.**
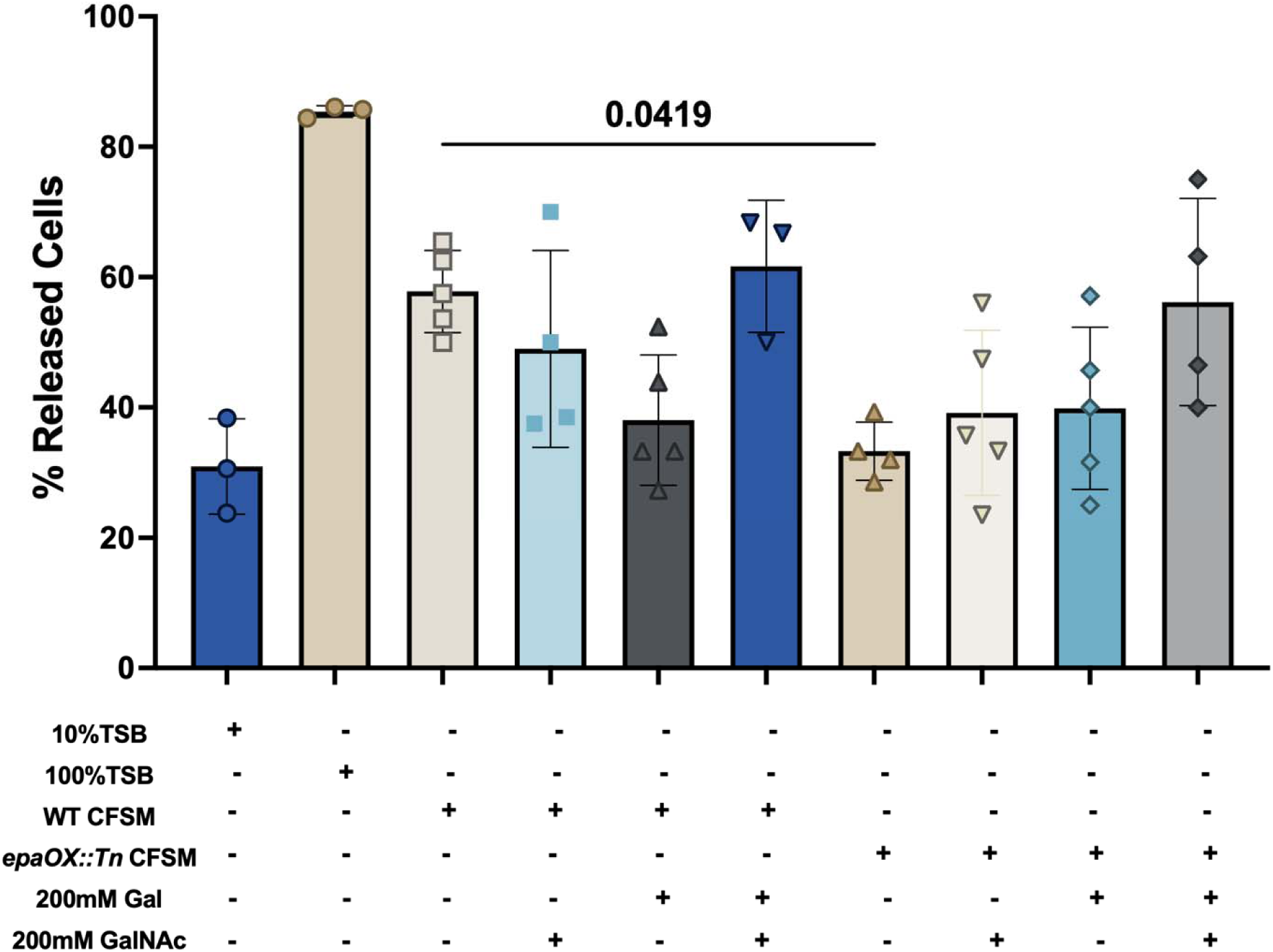
Galactose and GalNAc restore stimulation of dispersion by *epaOX::Tn* CFS. Cell-free supernatant (CFS) from wild-type OG1RF and *epaOX::Tn* cultures were supplemented with 200 mM galactose (Gal) and/or GalNAc and applied to 4-day wild-type biofilms for 1 hour followed by harvest of the bulk liquid and attached biofilm and quantification of CFUs. Data are presented as percentage of released cells. Data are combined from 3–5 independent experiments. Statistics: Kruskal-Wallace one-way ANOVA followed by Dunn’s test versus *epaOX::Tn* CFS.

**Figure 7.**
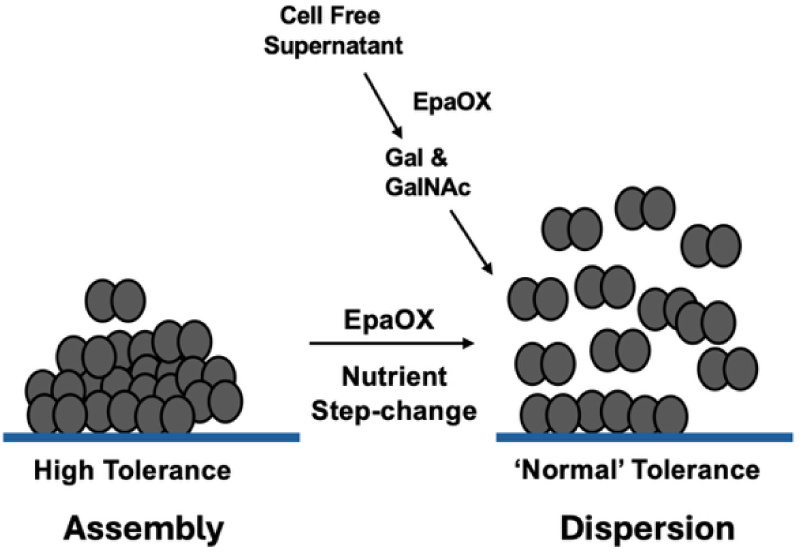
Model: Nutrient step change induces biofilm dispersion and disassembly in a manner that requires EpaOX within the biofilm. Nutrient step-change induced dispersion quickly eliminated the high phenotypic tolerance to antibiotics. Reversal of antibiotics tolerance was independent of mechanism of action of the drug. Dispersed cells maintained a high capacity for adherence. Cell free supernatant also induces biofilm dispersion, suggesting that cell-to-cell interactions play a role in triggering dispersion. EpaOX was also necessary for cell-free supernatant induced dispersion and supplementation of *epaOX::Tn* CFS with galactose and GalNAc was sufficient to restore CFS mediated dispersion to wild type levels.

## Discussion

Biofilms are dynamic microbial systems in which cells switch between attached and planktonic states. Mature biofilms are constantly changing in structure, which can ultimately result in the detachment of certain populations of cells and partial collapse of biofilm architecture (Allison et al. 1990; Karin Sauer et al. 2002). In *E. faecalis* most biofilm studies have focused on assembly and defining the pathway of biofilm assembly. This has resulted in the production of a multi-stage model of biofilm formation (Ch’ng et al. 2019; Willett and Dunny 2024) similar to the developmental model proposed in other systems (O’Toole, Kaplan, and Kolter 2000). We present here data that suggests *E. faecalis* biofilm dispersion occurs similar to well-described systems such as *P. aeruginosa,* with fast kinetics in response to both nutrient availability and cell-to-cell signaling. Importantly, triggering dispersion also results in a fast and significant reduction of the high phenotypic tolerance to antibiotics that makes *E. faecalis* biofilm infections difficult to treat.

We began to describe dispersion using a model of a sudden increase in nutrients (Fig 1). Nutrient step-change induced dispersion in our system as measured by the number of released cells versus remaining attached cells (Fig. 1) and by confocal microscopy (Fig. 2). The series of events for individual cells within a biofilm is, presumably, to first sense and transduce a signal of increased nutrient availability. In *Firmicutes*, nutrient-responsive gene expression is regulated by *ccpA*, *codY*, and the stringent response (Gaca and Lemos 2019). The *ccpA* and *codY* transcription factors regulate gene expression in response to glucose and GTP levels, respectively. In *E. faecalis* both *ccpA* (Gao et al. 2013) and *codY* (Colomer-Winter et al. 2019) were necessary for biofilm formation. Mutants in *relA* and *relQ*, which have reduced cellular concentrations of the stringent response alarmone (p)ppGpp, exhibited reductions in biofilm biomass and survival after the initial attachment stage (Colomer-Winter et al. 2019). These data are consistent with dispersion, suggesting that (p)ppGpp may actively maintain *E. faecalis* biofilms. In response to a sudden increase in nutrients, CodY-dependent promoters are likely to be repressed, resulting in reduced levels of (p)ppGpp, which may trigger biofilm dispersion. In *P. aeruginosa*, induction of the stringent response and (p)ppGpp was sufficient to induce both biofilm density and antibiotic tolerance in a dose-dependent manner (Engelhardt et al. 2025).

We also described the induction of dispersion by cell-free supernatants of stationary phase OG1RF liquid cultures (Fig 4). These data suggest that *E. faecalis* responds to soluble molecules released into the local environment in the decision to disassemble biofilms. The quorum-sensing system Fsr is necessary for biofilm assembly in vitro (Pillai et al. 2004; Hancock and Perego 2004) but also may be necessary for long-term maintenance of biofilms (Ali, Lévesque, and Neelakantan 2022). Recently, Fsr was found to be repressed during fluid flow and as a suppressor of late-stage biofilm growth in infective endocarditis (Antypas et al. 2026). The quorum signaling molecule AI-2 also was found to be sufficient to induce disassembly of established *E. faecalis* biofilms, but this was accomplished by the induction of prophages (Rossmann et al. 2015). These data suggest that quorum signaling may mediate transitions between adherence and dispersion in *E. faecalis* biofilms.

We found that dispersion induced by cell-free supernatant treated with heat and Proteinase K had decreased stimulation of dispersion (Fig 4), suggesting that secreted proteins may play a role. In some organisms, glycoside hydrolases have the potential to break down the exopolysaccharide matrix and release cells from the biofilm (Rumbaugh and Sauer 2020). OG1RF, for example, encodes 11 predicted glycoside hydrolases, so it is possible that a disassembly inducing enzyme remains to be discovered, although glycoside hydrolases of family GH18 purified from *E. faecalis* V583 did not disperse biofilms of *Staphylococcus epidermidis* (Keffeler et al. 2021). Other secreted proteins that affect the extracellular DNA necessary for cell attachment *in vitro*, or hydrolases that break down cell envelope components such as wall teichoic acid and peptidoglycan, may also mediate the release of cells from established biofilms.

We described a role for the glycosyltransferase *epaOX* in stimulating biofilm dispersion in CFS. EpaOX is predicted to contribute to ‘decoration’ of the rhamnose backbone of Epa polysaccharides with repeating subunits rich in galactose and GalNAc (Guerardel et al. 2020; Davis et al. 2025). Previous mass spectrometry in *epaOX* deletion mutants also showed reduced levels of free galactose and GalNAc (Dale et al. 2017). Our extracts of cell surface carbohydrates did not promote dispersion (Fig S6). However, we found that galactose and GalNAc added to *epaOX::Tn* CFS were sufficient to at least partially restore dispersion of wild-type biofilms (Fig 6). It is possible that the supplemented galactose and GalNac were enzymatically added to Epa polysaccharides in the CFS. Presumably, some Epa substrate as well as secreted glycotransferases would survive the filtering step. Dispersed cells of the *epaOX::Tn* strain had a reduced capacity to re-attach to multi-well plates compared to WT dispersed cells (Fig 5C). Other mutations in the *epa* locus have also been shown to be attenuated in *Caenorhabditis elegans* infection (Ocvirk et al. 2015), zebrafish infection (Prajsnar et al. 2013; Smith et al. 2019), a mouse model of peritonitis (Teng et al. 2009; Xu et al. 2000) and catheter-induced ascending urinary tract infection (Singh, Lewis, and Murray 2009). It is possible that reattachment of dispersed biofilm cells may play a key role in perpetuating systemic infections.

Dispersed cells of *P. aeruginosa* have altered expression for genes that regulate motility, metabolism, and surface attachment when compared to planktonic cells and cells within a biofilm (Kalia and Sauer 2024). Released OG1RF cells had an increased capacity for attachment to a surface when compared to stationary phase or exponential phase planktonic cells (Fig 1). This suggests dispersed cells are poised to rapidly re-establish on a surface as soon as they are released from the biofilm. This would facilitate re-attachment and the formation of new microcolonies at distant locations. Dispersed cells, however, also quickly lose their increased phenotypic tolerance to antibiotics that is characteristic of biofilms (Fig 3). Loss of tolerance occurred for all tested antibiotics regardless of their mechanism of action.

In conclusion, our study has shown that *E. faecalis* biofilms dynamically respond to changes in their environment, especially to changes in nutrients. Nutrient-induced dispersion rapidly reverses the phenotypic high antibiotic tolerance in dispersed biofilm cells. Certain molecules produced during the culture of *E. faecalis* also appear to stimulate dispersion. Triggering dispersion has the very real possibility of also inducing systemic infection and septic shock (Fleming and Rumbaugh 2018). However, it may be possible to use dispersion genetics to design adjuvants that increase the effectiveness of antibiotics against biofilms. This would require dispersion to be described in sufficient molecular detail to genetically uncouple the reversal of biofilm phenotypic antibiotic tolerance from the mechanical disassembly of the biofilm itself.

## Acknowledgements

We thank members of the Binghamton Biofilm Research Center, especially Manmohit Kalia, Soyoung Park and Karin Sauer for helpful discussions. This work was supported by startup funds from Binghamton University and Albany Medical College, Research Experiences for Undergraduates (REU) in Microbial Biofilm Development, Resistance, & Community Structure (NSF, 2150229 and 2349311 Caitlin Light and Karin Sauer, PIs), Bridges to the Baccalaureate Research Training Program (T34GM154616, Lisa Savage, PI), a grant from Small Scale Systems Integration and Packaging (S3IP) Center of Excellence, funded by New York Empire State Development Division of Science, Technology and Innovation to PTM, Bridges Student Fellowship to MK, McNair Program Summer Fellowship to RM-A, Binghamton University External Scholarships and Undergraduate Research Center Undergraduate Research Awards to RM-A and MK and the Binghamton University Graduate School Doctoral Focus Fellowship to NM.

## Author Contributions

NM Performed all experiments except Figure S1 and S2 performed by DML. RM-A and MK assisted with some experiments. LCC, DGD and PTM conceived and designed the study. NM and PTM wrote the paper with inputs from all the co-authors.

---

**Supplementary Figure 1:**
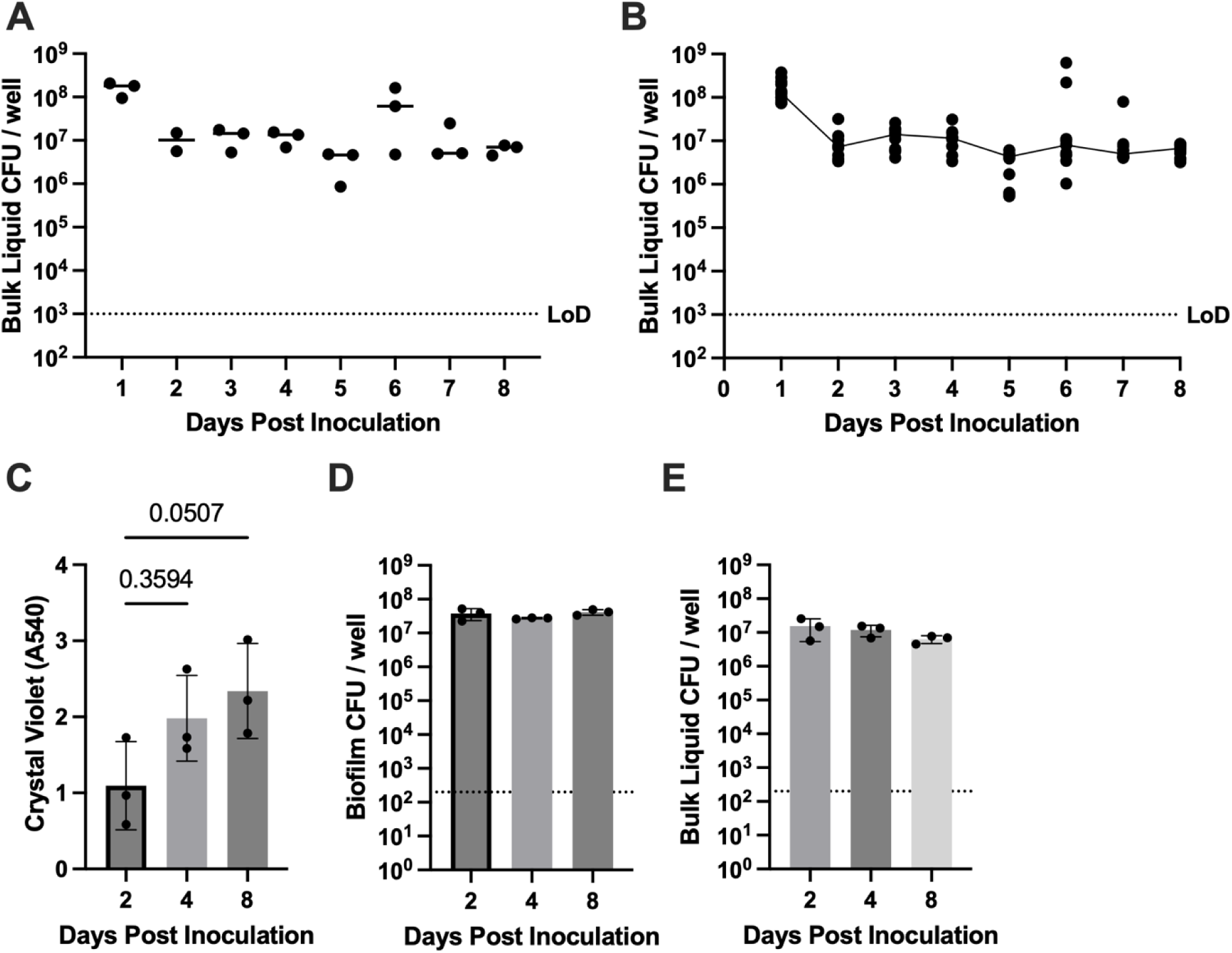
OG1RF forms stable long-term biofilms and does not undergo spontaneous dispersion in these conditions. A) CFU content of the bulk liquid of long-term biofilms after overnight growth. Each data point represents the mean of 3-4 individual wells from 3 independent experiments. B) CFU content of each technical replicate well from A. The apparent increase in mean CFUs at day 6 was caused by two outlier wells. C) Crystal Violet staining of biofilm biomass over 8 days with daily medium change. Each data point represents the mean of 3-4 individual wells from 3 independent experiments. Statistics: Kruskal-Wallace One-way ANOVA with Dunn’s test versus Day 2. D) Attached biofilm CFUs over 8 days, E) Bulk Liquid CFUs over 8 days. Data from D and E: 3 independent experiments, each data point represents the mean of 3-4 technical replicate wells. Statistics: Kruskal-Wallace One-way ANOVA with Dunn’s test versus Day 2

**Supplementary Figure 2:**
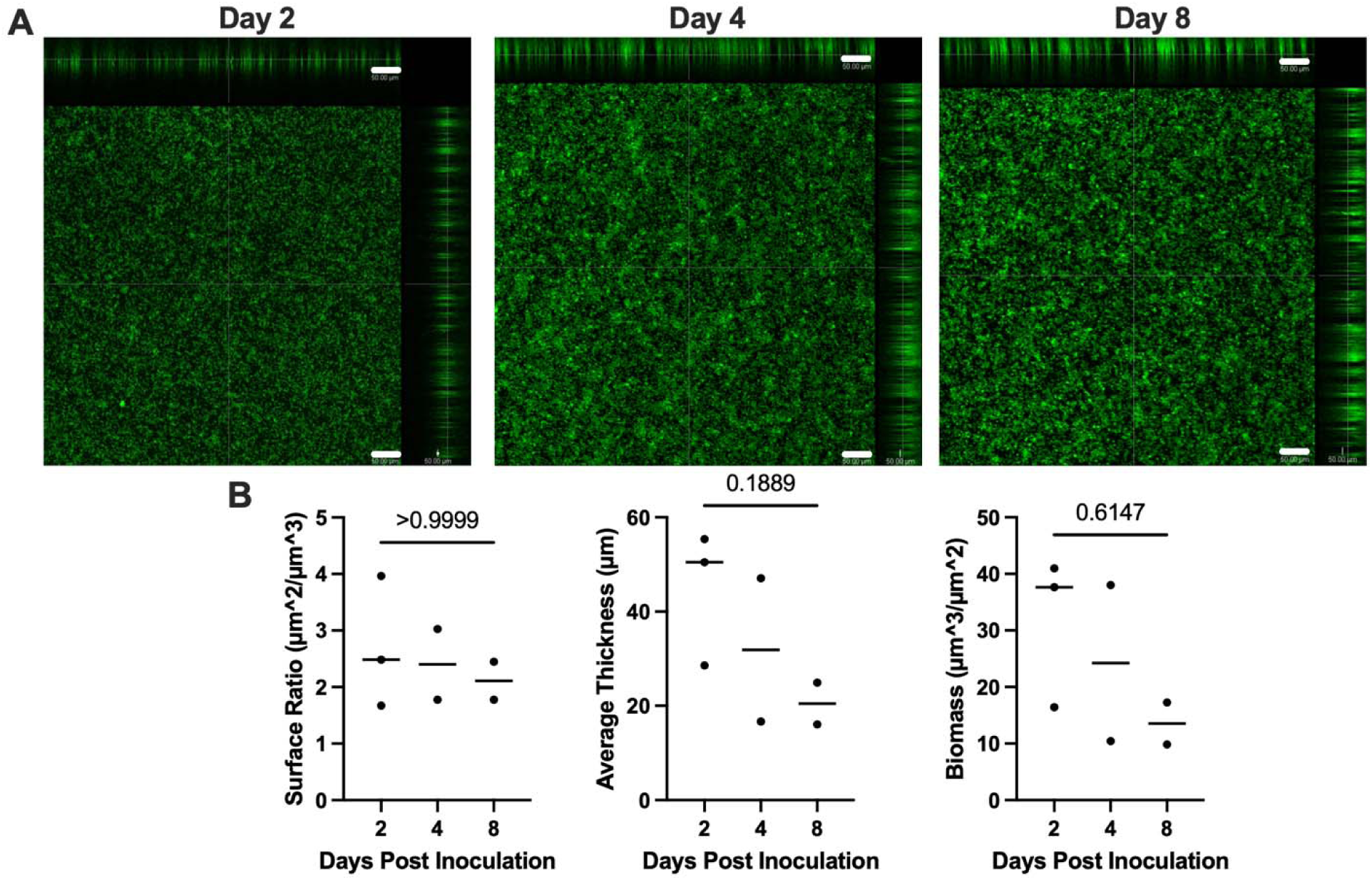
Confocal Laser Scanning Microscopy of long-term biofilms. A) Representative field z-stack projections of wild type OG1RF biofilms stained with SYTO-9 at days 2, 4 and 8 post inoculation. Magnification = 10x, Scale Bar = 50μm. B) Quantification of biofilm microscopy with COMSTAT-2, each data point represents the mean of 2 fields acquired from four technical replicate wells.

**Supplementary Figure 3:**
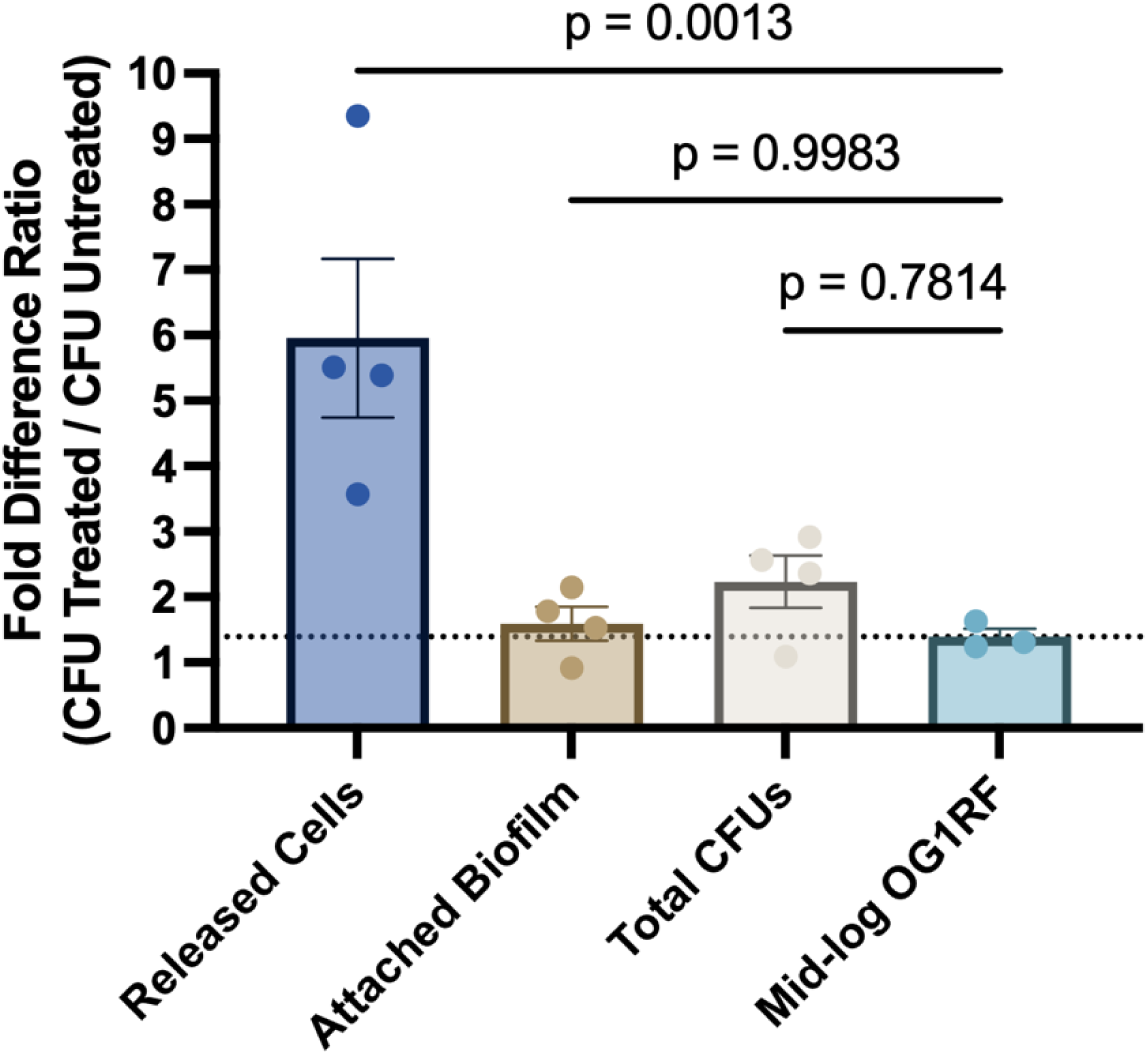
**Cell release does not represent cell growth**: Day 4 biofilms of OG1RF grown in 10% TSB with media changes every 24-hours were washed in fresh 10% TSB, then 100% TSB was added for 1 hour. 10% TSB was added to untreated control wells. ‘Released’ CFUs were quantified by plating the overlying liquid. ‘Attached’ CFUs were quantified by mechanical disruption and plating of remaining biofilm. ’Total’ CFUs = ‘Released Cells’ + ‘Attached Biofilm’. Data were compared to a mid-log liquid culture of OG1RF grown in 10% TSB that was shifted to 100% TSB, grown for 1 hour at 37C then plated. This represents the maximum possible growth of OG1RF under assay conditions. Data are presented as fold difference ratio of Treated CFUs / Untreated CFUs. Each data point represents the mean of 3-4 technical replicate wells combined from 4 independent experiments.

**Supplementary Figure 4.**
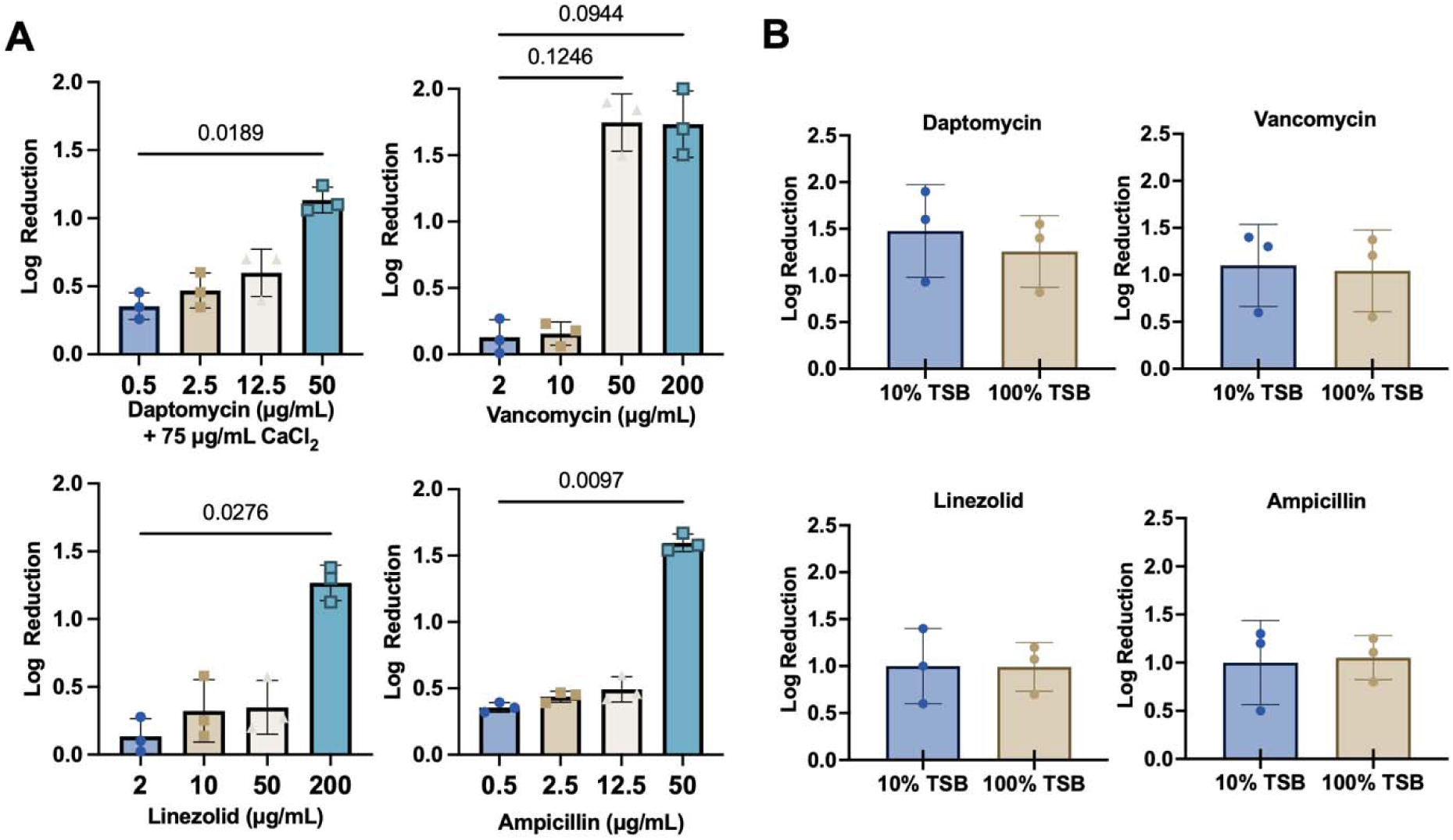
Antibiotic dose determination and nutrient-dependent antibiotic susceptibility in OG1RF. **(A**) Mid-log OG1RF grown in 10% TSB was exposed to the indicated concentration of antibiotic for 1 hour followed by washout and plating. Data represent 3 independent experiments, each with 3–4 technical replicate wells (n = 3 biological replicates). Statistics: Kruskal-Wallace One-way ANOVA with Dunn’s test versus the lowest concentration. **(B)** Effect of nutrient conditions on antibiotic susceptibility. Mid-log OG1RF were treated with the selected antibiotic concentrations in either low-nutrient (10% TSB) or high-nutrient (100% TSB) medium, followed by wash out and plating to evaluate the influence of nutrient availability on antibiotic efficacy. Daptomycin = 50μg/mL + 75μg / mL CaCl_2,_ Vancomycin = 50μg/mL, Linezolid = 200μg/mL, Ampicillin = 50 μg/mL Data represent 3 independent experiments, each with 3–4 technical replicate wells. Statistics: Unpaired Welch’s t-test, all comparisons were not significant.

**Supplementary Figure 5.**
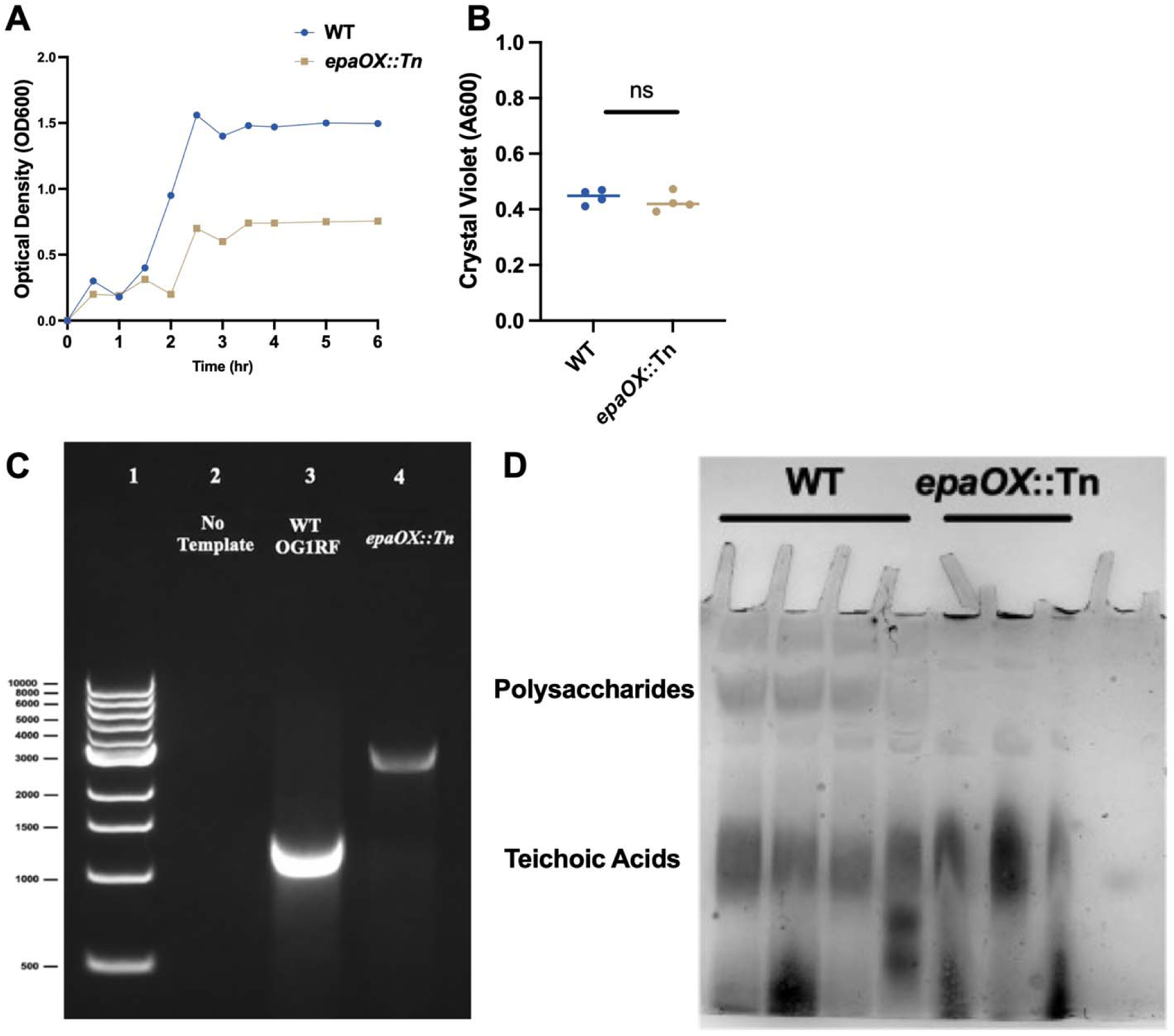
Characterization of *epaOX::Tn*. **(A)** Growth kinetics of OG1RF WT and *epaOX::Tn* of the indicated culture in 100% TSB by OD600. **(B)** Biofilm biomass quantification. Four-day mature biofilms of WT and *epaOX::Tn* were assessed using crystal violet staining in 24-well plates. Data represent 4 independent experiments, each with 3–4 technical replicate wells. Statistics: unpaired Welch’s t-test **(C)** Agarose gel electrophoresis of PCR genotyping of *epaOX::Tn* with primers that flank *epaOX.* The results confirm the presence of the 2kb transposon within *epaOX*. Lane 1 – 1kb ladder, Lane 2 No template control, Lane 3 WT OG1RF, Lane 4 *epaOX::Tn*.

**Supplementary Figure 6.**
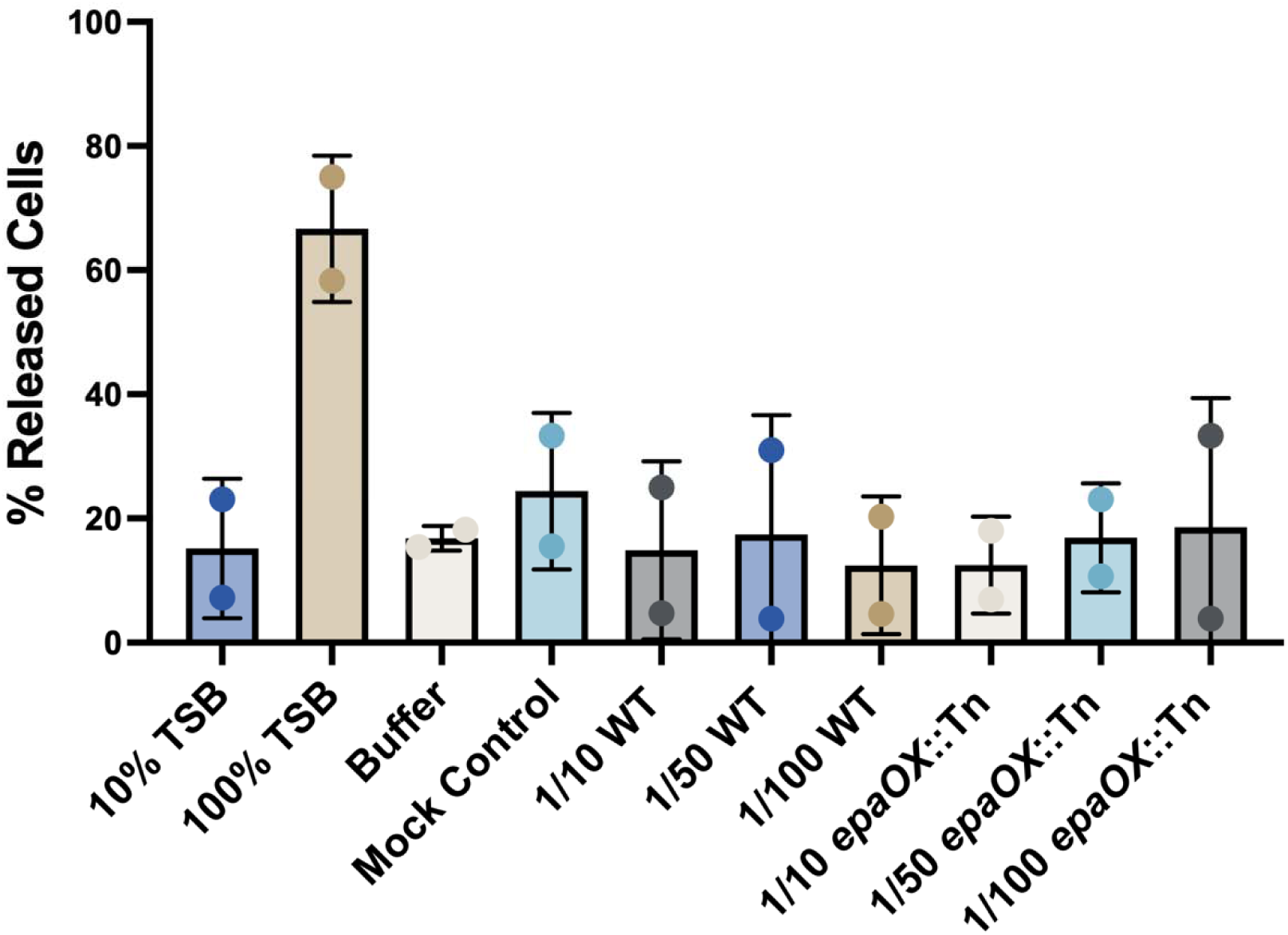
Carbohydrate-enriched extracts are not sufficient to stimulate biofilm dispersion. Carbohydrate-enriched extracts isolated from OG1RF WT and *epaOX::Tn* were applied to 4-day mature WT biofilms for 1 hour and the percentage of released cells was compared to untreated controls. Carbohydrate-enriched extracts (final volume 50 µL) were applied at fractional inputs corresponding to 1/10, 1/50, and 1/100 of the total preparation in 10%TSB. Data represent 2 independent experiments, each with 3–4 technical replicate wells (n=2 biological replicates).

**Table S1:**
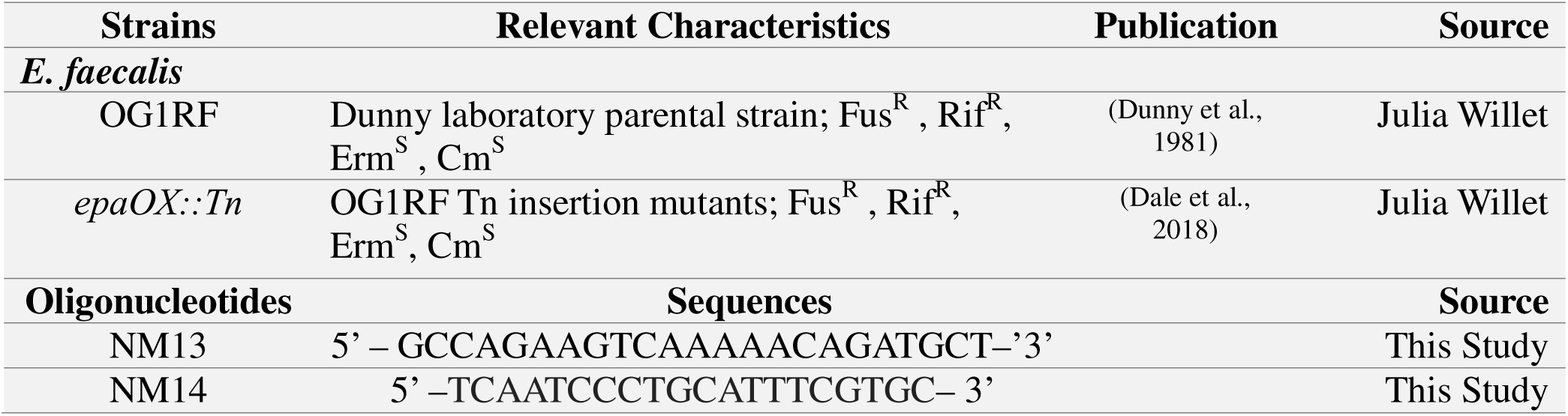
Strains and primers used in this study.

## Methods

### Bacterial strains and growth conditions

*Enterococcus faecalis* OG1RF is a Rifampicin and Fusidic acid resistant derivative of E. faecalis OG1, which was first isolated from a human oral cavity (Dunny et al., 1978). For *Enterococcus faecalis* OG1RF wild-type strain, it was maintained as freezer stocks at −80°C in 50% (v/v) glycerol solution. The OG1RF strain was routinely grown in Tryptic Soy Broth (BD) for generating freezer stocks. OG1RF strain was streaked from stocks before each replicate experiment on Enterococcus Selective Agar. Bacterial overnight cultures were grown in the same medium used for experiments in flasks with continuous shaking at 180 rpm until mid-log (OD 0.5) measured by OD600 Spectrophotometer at 37°C in an aerobic incubator. Antibiotics were used at the following concentrations: Rifampicin (RIF), 50μg/ml and Fusidic acid (FA), 25μg/ml for OG1RF growth and 10μg/ml of Chloramphenicol (CHL) for epaOX::Tn growth.

### Biofilm Growth

Biofilms were grown in semi-continuous culture systems in 24-well plates with each well having a growth area of 1.9 cm^2^, allowing for sufficient biofilm biomass development for microscopy and antibiotic tolerance assays. Using a mid-log overnight (OD=0.5), an inoculum was made by diluting 1:10 into a 10-fold diluted TSB medium. 500μL of inoculum was transferred into each well of 24-well treated tissue culture plates (Genclone, Genesee Scientific). 500 μL of sterile 10% TSB was added to the bottom row wells, acting as contamination checks and to prevent evaporation of samples. Biofilms were grown stationary at 37°C for a time-course of four days. Media changes were performed every 24h by removing the spent media slowly to prevent disruption of biofilm and replacing spent media with fresh 500 μL of 10% TSB.

### Native Dispersion

Biofilm were grown in a time course of 5-8 days, and every day, biofilms were harvested by dispensing 500μL of sterile PBS into each well and scraping vigorously in a zigzag pattern. Scraping was repeated twice per well to ensure maximum yield. Samples were then homogenized for 10 seconds using an OMNI TH tissue homogenizer with a one mL probe. Samples were then serially diluted and plated in Tryptic Soy agar incubated at 37 °C in a stationary position for 24 hours. Native Dispersion events were indicated by an increase in CFUs/ml.

### Nutrient-induced Dispersion Assays in 24-well Plates

Following biofilm growth for a total of four days with medium change every 24 hours, the spent medium was removed from each well and replaced with 500 μL of fresh 10-fold diluted TSB and incubated stationary at 37°C for 1h, serving as a wash step prior to treatment to remove any loosely adherent cells. After one hour, the wash liquid was removed and discarded and replaced with 500 μL control (10% TSB) or treatment (100% TSB) and incubated stationary for 1 h at 37°C. Released cells in the overlaying bulk liquid were collected into microfuge tubes with 500 μL of sterile PBS. The remaining biofilm was harvested from the bottom of the well by dispensing 500μL of sterile PBS into each well and scraping vigorously in a zigzag pattern, repeated twice per well. Samples were then homogenized for 10 seconds using an OMNI TH tissue homogenizer with a one mL probe. Homogenized samples were then 10-fold serially diluted and drop-plated onto Tryptic Soy Agar (TSA) and incubated at 37°C. ‘Released’ CFUs were quantified by plating the overlying bulk liquid. ‘Attached’ CFUs were quantified by mechanical disruption and plating of remaining biofilm. ’Total’ CFUs = ‘Released Cells’ + ‘Attached Biofilm’.

### Antibiotic susceptibility assay for planktonic cells

To determine a dose of each antibiotic that reduced CFUs of exponentially growing OG1RF by 1-2 logs after 1 hour of exposure to the drug, OG1RF (OD=0.5) was grown to mid-log phase in 10 and 100% TSB, then 1ml of the sample was distributed into a microfuge tube. Samples were then treated with either different concentrations of the antibiotics based on the minimum inhibitory concentration (MIC) value determined from (Frank et al. 2015) or untreated with the addition of sterile distilled water. Antibiotic concentrations ranged from MIC to 100 x MIC. All Daptomycin assays were supplemented with 75μg/mL of CaCl2 to maintain physiological calcium levels (Fuchs, Barry, and Brown 2000). The samples were incubated stationary for 1h. Antibiotics were removed by centrifugation at 8000rpm for 1 minute, then carefully removing the supernatant and replacing it with 1mL of sterile PBS. The samples were then 10-fold serially diluted and drop-plated in Tryptic Soy Agar.

### Antibiotic susceptibility assay for attached and released biofilm cells

Biofilms were grown in 24-well plates for 4 days with medium change every 24 hours. They were washed with 10% TSB for 1 hour and then were exposed to 100% TSB or 10% TSB for 1hour. For each Bulk liquid sample, it was removed from the wells and vortexed, then distributed equally to the following treatments: Ampicillin (VWR Life Science) at a concentration of 50 µg/ml, Linezolid (Chem-Impex) at a concentration of 200 µg/ml, Vancomycin (Chem-impex) at a concentration of 50 µg/ml, Daptomycin (Combi-blocks) at a concentration of 50 µg/ml + 75 µg/ml CaCl_2_ then incubated stationary at 37°C for 1 hour. After 1h, bulk liquid samples were centrifuged at 8000rpm for 1 minute, and the supernatants were carefully removed and replaced with 1mL of sterile PBS. For Biofilm samples, 500 μL of PBS with the appropriate concentration of the antibiotic used was added to treated wells and 500 μL sterile PBS to untreated wells, then incubated stationary at 37°C for 1h. For biofilm samples, the treatment was removed and was harvested by scraping into 500μL sterile PBS. Samples were serially diluted and drop-plated into Tryptic Soy Agar.

### Analysis of the biofilm structure

#### CLSM

Following 4-day biofilm growth in 24-well plates with medium change every 24 hours under static conditions, spent media were gently removed, and biofilms were washed once with sterile phosphate-buffered saline (PBS). Biofilms were then stained using the LIVE/DEAD BacLight Bacterial Viability Kit (Invitrogen, Carlsbad, CA) for 15 minutes at room temperature, according to the manufacturer’s instructions. Following staining, biofilms were fixed with 4% paraformaldehyde (PFA) for at least 20 minutes before imaging. Confocal laser scanning microscopy (CLSM) was performed using a Zeiss LSM 880 confocal microscope (Carl Zeiss Microscopy, Oberkochen, Germany) equipped with an LD Plan-Neofluar 10×/korr M27 objective lens. Image acquisition was conducted using ZEN Black (version 2.3) under the following parameters: scan speed of 7, averaging of 2×, pinhole size set to 1 Airy Unit, and 16-bit image depth. Imaging was performed at room temperature. Fluorophores were excited using an Argon laser (458, 488, and 514 nm) for SYTO-9 and a DPSS 561 nm laser for propidium iodide (PI). Emission detection ranges were set to 493–578 nm for SYTO-9 and 578–735 nm for PI. Z-stack images were acquired over a total depth of 30 µm, consisting of 20 optical slices with a step size of 1.5 µm. Sequential (frame-by-frame) acquisition was used to minimize spectral overlap between channels. It is important to note that for downstream analysis, the green fluorescence channel (SYTO-9) corresponds to Channel 1 in ZEN software but is exported as Channel 2 in the raw OME-TIFF files, and this mapping was accounted for during analysis. Raw image files were converted to OME-TIFF format using the Bio-Formats plugin in ImageJ, ensuring compatibility with downstream analysis tools. Quantitative analysis of biofilm biomass and surface area was performed using COMSTAT 2.1 (Heydorn et al. 2000), which calculates biomass (µm³/µm²) from three-dimensional image stacks. Biofilms were grown under two conditions: control (10% TSB) and treatment (100% TSB). For each condition, twelve independent image stacks were acquired and analyzed per replicate, across a total of four independent experiments.

#### Crystal violet staining

Biofilms were grown as before for four days with medium change every 24 hours. The spent medium was removed from each well followed by wash with 200 μL of PBS. Biofilms were fixed with 200 μL of 4% PFA for 10 minutes. Then, PFA was removed from each well and washed with PBS. 200 μL of 0.1% crystal violet was added to each well and allowed to stain for 10 minutes. Crystal violet was removed from the wells, and each well was washed with 200 μL PBS 5-6 times until the liquid became relatively clear. Stained biofilms were dried for about 20 minutes. Crystal violet was eluted with 200 μL of 95% ethanol and transferred into a 96-well plate along with an eluted blank well. 96-well plates were read at OD 570 (Genesys 30 Visible Spectrophotometer, Thermo Scientific).

### Attachment assay (2h) and (24h) for overall biomass

Dispersed cells were collected following nutrient-induced dispersion with 100% TSB as described above. The dispersed cells were harvested from the supernatant and the optical density adjusted to OD = 0.2 in fresh 10% TSB. For comparison, *E. faecalis* exponential-phase (OD_600_ < 0.5) cultures were grown in 10% TSB at 37°C with shaking to mid-log phase. Stationary-phase cultures (OD_600_ > 1.3) were grown for 16–18 h under the same conditions. Both exponential and stationary-phase cells were similarly adjusted to OD = 0.2 in 10% TSB. Aliquots (100 µL) of each cell added to individual wells of tissue culture treated 96-well plates. Blank wells containing only 10% TSB served as negative controls. To minimize edge effects due to evaporation, sterile water was added to all unused wells surrounding the samples. Plates were incubated at 37°C under static conditions for either 2 h (early attachment) or 24 h (biofilm maturation), with a single media replacement after 12 h in the 24-h condition. Each treatment was performed using 4–8 technical replicates. After incubation, non-adherent cells were removed by carefully aspirating the supernatant. Wells were washed 3–4 times with 200 µL PBS and stained with 0.1% crystal violet solution and incubated for 15 min at 37°C. Excess stain was removed by washing the wells twice with distilled water. Plates were then air-dried upside down on tissue paper for 20 min. For qualitative assessment, plates were photographed after drying. Biofilm-bound CV was solubilized with 200 µL of 95% ethanol per well and incubated for 15 min at room temperature. The contents were briefly mixed by pipetting, and 125 µL of the solubilized CV was transferred to a new plate. Absorbance was measured at 600nm using a microplate reader (Cerillo) to quantify biofilm biomass.

### Treatment of Cell-Free Spent Medium

Cell-free spent medium (CFS) was collected from 72 h OG1RF cultures by centrifugation and filtering with a 0.2 μm syringe filter. For heat treatment, WT CFS was incubated at 95°C for 30 min, cooled to 37°C, and applied to 4-day-old WT biofilms following the first medium replacement with 10% TSB. CFS samples were treated with proteinase K (Fisher Bioreagents) at final concentrations of 100µg/ml and incubated at 60°C for 1 hour followed by heat inactivation at 80°C for 30 min. Treated supernatants were stored at 4°C until use and applied to biofilms after the first medium change with 10% TSB.

### Small-Scale Capsule Purification and Carbohydrate PAGE Analysis

Capsular polysaccharides were isolated using a modified small-scale extraction protocol as previously described ((Hancock and Gilmore 2002; Thurlow, Thomas, and Hancock 2009). Briefly, OG1RF Enterococcus faecalis were grown overnight in 2 mL tryptic soy broth (TSB) supplemented with 1% glucose with fusidic acid and rifampicin. Overnight cultures were diluted 1:100 into 25 mL fresh TSB containing 1% glucose and grown at 37°C to mid-log phase (OD = 0.6–0.8). Cells were harvested by centrifugation at 4,000 × g for 10 min at 4°C and resuspended in 2 mL capsule preparation buffer (25% sucrose, 10 mM Tris-HCl [pH 8.0], 0.05% sodium azide). Suspensions were divided into two microcentrifuge tubes and pelleted at 13,000 × g for 5 min at 4°C. Combined pellets were resuspended in 750 µL CPB and treated with lysozyme (1 mg mL ¹ final concentration) and mutanolysin (10 U mL ¹ final concentration) to digest the cell wall. Samples were incubated at 37°C for 16 h with gentle rotation. Cellular debris was removed by centrifugation at 16,000 × g for 20 min at 4°C, and the supernatant was treated with RNase (100 µg mL ¹), DNase I (100 U mL ¹), MgCl (2.5 mM), and CaCl (0.5 mM).

Samples were incubated at 37°C for 4 h with gentle rotation, followed by protein digestion with pronase (50 µg mL ¹ final concentration) for an additional 16 h at 37°C. Following enzymatic digestion, samples were centrifuged at 16,000 × g for 20 min at 4°C, and the supernatants were transferred to fresh tubes. Chloroform extraction was performed by adding 500 µL chloroform, mixing by inversion, and centrifuging at 16,000 × g for 5 min. The aqueous phase was recovered, and carbohydrates were precipitated by the addition of ethanol to a final concentration of 75% (v/v). Samples were incubated at −80°C for 30 min and centrifuged at 16,000 × g for 20 min to pellet polysaccharides. Pellets were washed once with 75% ethanol, air-dried at room temperature, and resuspended in 50 µL Tris-NaCl buffer (50 mM Tris-HCl, 150 mM NaCl, pH 8.0). Capsular polysaccharides were analyzed by carbohydrate polyacrylamide gel electrophoresis. Gels were stained overnight at room temperature in Stains-All solution (0.005% in 50% ethanol) in the dark and destained with destaining solution (50% methanol, 10 % acetic acid, and 40% water). Gels were visualized using ChemIDoc (BioRad). Capsule extracts were tested by addition to 10% TSB for dispersion testing of 4-day old biofilms as described above.

### Genotyping of *epaOX::Tn*

Genomic DNA was extracted from wild-type Enterococcus faecalis strain OG1RF and the epaOX::Tn mutant using the Wizard Genomic DNA Purification Kit (Promega) following the manufacturer’s protocol. DNA concentration and purity were assessed using a SpectraDrop™ Micro-Volume Microplate. To confirm the transposon insertion, PCR was performed using primers: NM13 and NM14, designed to flank the epaOX locus (including the full ORF plus ∼200 bp of upstream and downstream flanking sequence).

